# O-GlcNAcylation of SAMHD1 Indicating a Link between Metabolic Reprogramming and Anti-HBV Immunity

**DOI:** 10.1101/2020.03.09.983338

**Authors:** Jie Hu, Qingzhu Gao, Yang Yang, Jie Xia, Wanjun Zhang, Yao Chen, Zhi Zhou, Lei Chang, Yuan Hu, Hui Zhou, Li Liang, Xiaosong Li, Quanxin Long, Kai Wang, Ailong Huang, Ni Tang

## Abstract

Viruses hijack the host cell machinery to promote viral replication; however, the mechanism by which metabolic reprogramming regulates innate antiviral immunity in the host remains elusive. Herein, we found that Hepatitis B virus (HBV) infection upregulates glucose transporter 1expression, promotes hexosamine biosynthesis pathway (HBP) activity, and enhances O-linked β-N-acetylglucosamine (O-GlcNAc) modification of downstream proteins. HBP-mediated O-GlcNAcylation positively regulates host antiviral response against HBV *in vitro* and *in vivo*. Mechanistically, O-GlcNAc transferase (OGT)-mediated O-GlcNAcylation of sterile alpha motif and histidine/aspartic acid domain-containing protein 1 (SAMHD1) on Ser93 stabilizes SAMHD1 and enhances its antiviral activity. In addition, O-GlcNAcylation of SAMHD1 promoted its antiviral activity against human immunodeficiency virus-1 *in vitro*. In conclusion, the results of our study reveal a link between HBP, O-GlcNAc modification, and innate antiviral immunity by targeting SAMHD1. Therefore, the results of this study demonstrate a strategy for the potential treatment of HBV infection by modulating HBP activity.

## Introduction

Immunometabolism is an emerging field that highlights the importance of specific metabolic pathways in immune regulation. Metabolic enzymes, such as glyceraldehyde 3-phosphate dehydrogenase and pyruvate kinase isozyme M2 can directly modulate immune cell activation (Chang *et al*, 2013; Palsson-McDermott *et al*, 2015). In addition to providing energy and building blocks for biosynthesis, metabolites have been shown to participate in epigenetic modification and signaling transduction. the glycolytic product lactate not only regulates gene expression by histone acetylation (Zhang *et al*, 2019a), but also acts as a suppressor of type I interferon signaling by interacting with the mitochondrial antiviral signaling protein MAVS (Zhang *et al*, 2019b). Itaconate—another important metabolite for immune function—downregulates type I interferon signaling during viral infection by promoting alkylation of Kelch-like ECH-associated protein 1 and activation of anti-inflammatory proteins, including nuclear factor erythroid 2-related factor 2 (Mills *et al*, 2018, 1; O’ Neill & Artyomov, 2019).

Viruses are obligate parasites that rely on the biosynthetic machinery of the host to complete their life cycle. They hijack the host cell machinery upon entry to fulfill their energetic and biosynthetic demands for viral replication. Human cytomegalovirus (HCMV) and herpes simplex virus-1 (HSV-1) remodel host cells to perform distinct, virus-specific metabolic programs (Vastag *et al*, 2011). HCMV reprograms host metabolism by upregulating the expression of carbohydrate-response element binding protein and glucose transporter 4 (GLUT4) to provide materials for viral replication(Yu *et al*, 2014).Glucose uptake, glycolysis, and lipogenesis are enhanced in HCMV-infected cells to synthesize biomolecules. Moreover, HSV-1 promotes central carbon metabolism to synthesize pyrimidine nucleotides.

On the other hand, hosts may recognize virus-induced signaling and reprogram metabolic pathways to protect themselves from further damage. Increased glucose utilization, increased aerobic glycolysis, and inhibition of oxidative metabolism have emerged as the hallmarks of macrophage activation (Jung *et al*, 2019). Pattern recognition molecules as well as several metabolic pathways and metabolites have been reported to play an important role in regulating host innate immune response (Haskó & Cronstein, 2004; Skelly *et al*, 2019; Tsalikis *et al*, 2013).Therefore, it is important to identify the key metabolites that regulate innate immune response during viral infection. Understanding the relationship between cell metabolism, innate immunity, and viral infection may provide insights to develop new therapeutic targets to control viral infection.

Recent studies have emphasized the emerging role of the hexosamine biosynthesis pathway (HBP)—a branch of glucose metabolism—in host innate immunity. HBP links cellular glucose, glutamine, acetyl-CoA, and uridine triphosphate (UTP) concentrations with signaling transduction(Hanover *et al*, 2012). Approximately 2–5% of the total glucose entering a cell is converted to uridine diphosphate N-acetylglucosamine (UDP-GlcNAc) (McClain & Crook, 1996)—the end-product of HBP—and serves as a donor for O-linked β-N-acetylglucosamine (O-GlcNAc) modification (also known as O-GlcNAcylation) (Torres & Hart, 1984). O-GlcNAc transferase (OGT) and O-GlcNAcase (OGA) are responsible for the addition and removal of N-acetylglucosamine (GlcNAc) from Ser and Thr residues of target proteins. Several key host proteins involved in immune modulation, including signal transducer and activator of transcription-3 (STAT3), MAVS, and receptor-interacting serine/threonine-protein kinase 3 (RIPK3), are targets for O-GlcNAcylation (Li *et al*, 2017, 2018, 2019a; Song *et al*, 2019). However, the mechanism by which HBP-mediated O-GlcNAc modifications enhance antiviral innate immunity remains to be fully understood.

Hepatitis B virus (HBV) infection causes liver diseases, including acute and chronic hepatitis, cirrhosis, and hepatocellular carcinoma, which is a major global public health concern (Tsai *et al*, 2018). Current therapies improve both the quality of life and survival of patients with hepatitis B. However, new therapeutic approaches are needed to achieve functional cure of HBV infection (Fanning *et al*, 2019).

In this study, we investigated metabolic responses of host cells to HBV infection. Our results show that HBP-mediated O-GlcNAcylation regulates the antiviral activity of SAMHD1. Moreover, OGT promotes O-GlcNAcylation on Ser93 to enhance SAMHD1 stability and tetramerization, which is important for its antiviral activity. Our study established a link between HBP, O-GlcNAc modification, and antiviral innate immunity by targeting SAMHD1, thereby providing a potential drug target for treating HBV and human immunodeficiency virus-1 (HIV-1) infection.

## Results

### HBV infection upregulates GLUT1 expression and enhances HBP activity and protein O-GlcNAcylation

To explore metabolic changes in response to HBV infection, a metabolomics assay was performed in AdHBV-1.3-infected HepG2 cells (HepG2-HBV1.3) and AdGFP-infected HepG2 cells (HepG2-GFP). Principal component analysis showed that HBV infection dramatically changes the intracellular metabolic profile of HepG2 cells (Fig. 1A). Several metabolic pathways, including central carbon metabolism, amino sugar and nucleotide sugar metabolism(Supplementary Fig.1A) were significantly affected. Recent studies have shown that glucose metabolism plays a key role in host antiviral immunity (Li *et al*, 2018; Song *et al*, 2019). Hence, we determined the effect of altering glucose metabolism in HepG2-HBV1.3 cells. The expression level of several intermediate metabolites in glucose metabolism, including 3-phospho-glycerate, GlcNAc, N-acetyl glucosamine 6-phosphate (GlcNAc-6-P), and UDP-GlcNAc-the end-product of HBP-was increased upon HBV infection (Fig. 1B-D). To confirm this result, we established a strain of HepG2 cells engineered to express the human solute carrier family 10 member 1 (*SLC10A1*, also called NTCP) gene (HepG2-NTCP cells), which allows them susceptible to HBV infection (Hu *et al*, 2019). Targeted liquid chromatography-tandem mass spectrometry (LC-MS/MS) results showed a significant increase in UDP-GlcNAc and glucose levels in HBV-infected HepG2-NTCP, stable HBV-expressing HepAD38 (a tetracycline (Tet) inducible HBV expression cell line) (Fig. 1E-F), and AdHBV-1.3-infected HepG2 (Supplementary Fig.1B-1C) cells. These results were consistent with those observed in HepG2.2.15, an HBV-replicating cell line (Li *et al*, 2015). Because OGT-mediated protein O-GlcNAcylation is highly dependent on the intracellular concentration of the donor substrate UDP-GlcNAc, we examined whether HBV infection can affect O-GlcNAc modification in host cells. Total protein O-GlcNAcylation in HBV-infected HepG2-NTCP cells significantly increased 6 to 9 days post HBV infection. A similar result was observed in HepAD38 (Tet-off) cells (3 to 7 days after Tet removal from the medium) (Fig. 1G). Further, GLUT1 expression was markedly enhanced in our HBV cell models (Fig. 1H-I and Supplementary Fig.1D-E). Elevated glucose levels can increase HBP flux and enhance UDP-GlcNAc synthesis (Housley *et al*, 2008). However, we did not observe significant changes in the protein levels of OGT, OGA, and GFPT1—the key enzymes that regulate HBP flux and protein O-GlcNAcylation (Supplementary Fig.1F-G).These findings demonstrate that HBV infection upregulates GLUT1 expression, promotes glucose uptake, and increases UDP-GlcNAc synthesis and protein O-GlcNAcylation in host cells.

**Fig. 1.**
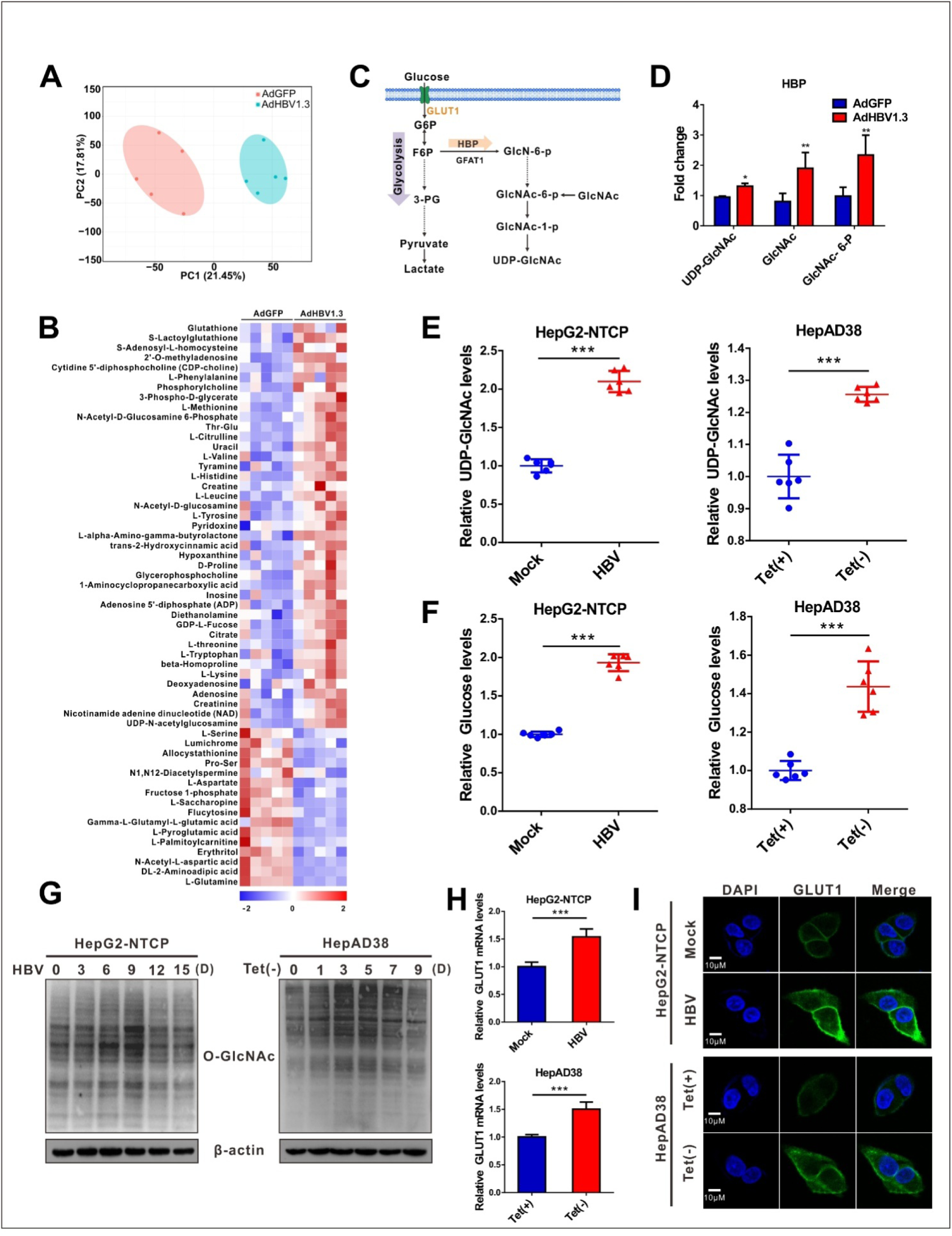
HBV infection promotes HBP and increases protein O-GlcNAcylation. (**A**) Principal component analysis of metabolite profiles obtained using a metabolomics assay in HepG2 cells infected with AdHBV1.3 or AdGFP for 72 h. (**B**) Heatmap of differentially expressed metabolites subjected to identical treatment conditions as in (a). n = 5. (**C**) An overview of the hexosamine biosynthesis pathway (HBP). (**D**) Fold changes in the expression of differentially expressed intermediate metabolites of HBP. n = 5. (**E**-**F**)Fold change in the expression of UDP-GlcNAc (E) and glucose (F) in HBV-infected HepG2-NTCP cells and HepAD38 cells with tetracycline inducible (Tet-off) HBV expression was determined using the LC-MS/MS targeted metabolomics assay. n = 6. (**G**) Immunoblot of total O-GlcNAc from HepG2-NTCP and HepAD38 cells treated for the indicated periods. (**H**-**I**) qPCR quantification (H) and immunofluorescence staining (I) of GLUT1 in HepG2-NTCP and HepAD38 cells, DAPI (blue) was used to counterstain nuclei, n = 9. Scale bar, 10 μm. Data are expressed as the mean ± SD. *P* values were derived from unpaired, two-tailed Student’s *t*-test in E, F, and H; (\*\*\**P*<0.001).

### Inhibition of protein O-GlcNAcylation promotes HBV replication in host cells

Next, we evaluated the effects of protein O-GlcNAcylation on HBV replication. HBV-infected HepG2-NTCP cells, HepAD38 (Tet-off) cells, and AdHBV-1.3-infected HepG2 cells were treated with inhibitors of GLUT1, GFPT1, OGT, and OGA. Pharmacological inhibition of GLUT1, GFPT1, and OGT reduced total protein O-GlcNAcylation levels (Fig. 2A-C, Supplementary Fig. 2A-C and Supplementary Fig. 3A-C), and promoted HBV replication (Fig. 2D-I, Supplementary Fig. 2D-F and Supplementary Fig. 3D-F). Conversely, pharmacological inhibition of OGA increased protein O-GlcNAcylation levels (Fig. 2J, Supplementary Fig. 2G and Supplementary Fig. 3G) but suppressed HBV replication (Fig. 2K-L, Supplementary Fig. 2H and Supplementary Fig. 3H). These data suggest that HBP-mediated O-GlcNAcylation positively regulates host antiviral immune response against HBV. The results of pharmacological inhibitor studies were similar to those obtained from shRNA-mediated knockdown of *GLUT1, GFPT, OGT*, or *OGA* in HepAD38 (Tet-off), HBV-infected HepG2-NTCP, and AdHBV-1.3-infected HepG2 cells (Fig. 3 and Supplementary Fig. 4). Taken together, these results indicate that inhibition of HBP or protein O-GlcNAcylation promotes HBV replication, whereas increased O-GlcNAc modifications can enhance host antiviral innate immune response against HBV.

**Fig. 2.**
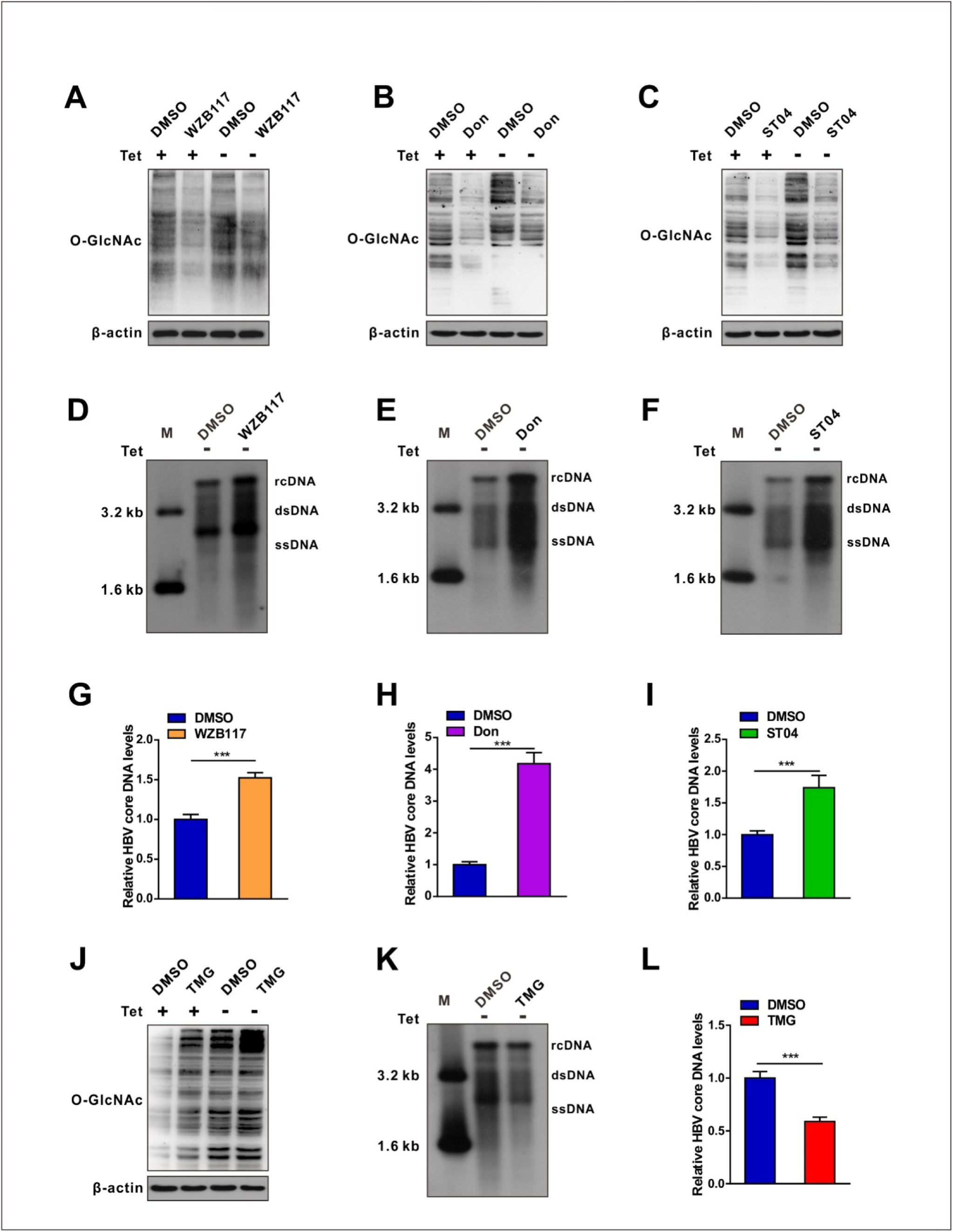
Pharmacological inhibition of protein O-GlcNAcylation promotes HBV replication. (**A-C**) Immunoblot of total O-GlcNAc from tetracycline-inducible HepAD38 cells treated with or without GLUT1 inhibitor WZB117 (50 μM) (A), GFPT1 inhibitor Don (30 μM) (B), or OGT inhibitor ST04 (100 μM) (C) for 72 h. Don, 6-Diazo-5-oxo-L-norleucine; ST04, ST045849. (**D**-**F**) HBV DNA were detected by Southern blot assay in stable HBV-expressing HepAD38 cells treated as above. rc DNA, relaxed circular DNA; ds DNA, double-stranded DNA; ss DNA, single-stranded DNA. (**G**-**I**) Quantification of HBV core DNA levels in stable HBV-expressing HepAD38 cells treated as indicated using qPCR, n=9. (**J**) Immunoblot of total O-GlcNAc from tetracycline-inducible HepAD38 cells treated with or without OGA inhibitor TMG (100 μM) for 72 h. TMG, Thiamet G. (**K**-**L**) Southern blot analysis of HBV DNA and qPCR quantification of HBV core DNA levels in stable HBV-expressing HepAD38 cells treated as in (J), n=9. Data are expressed as the mean ± SD. *P* values were derived from unpaired, two-tailed Student’s *t*-test in G-I and L; (\*\*\**P*<0.001).

**Fig. 3.**
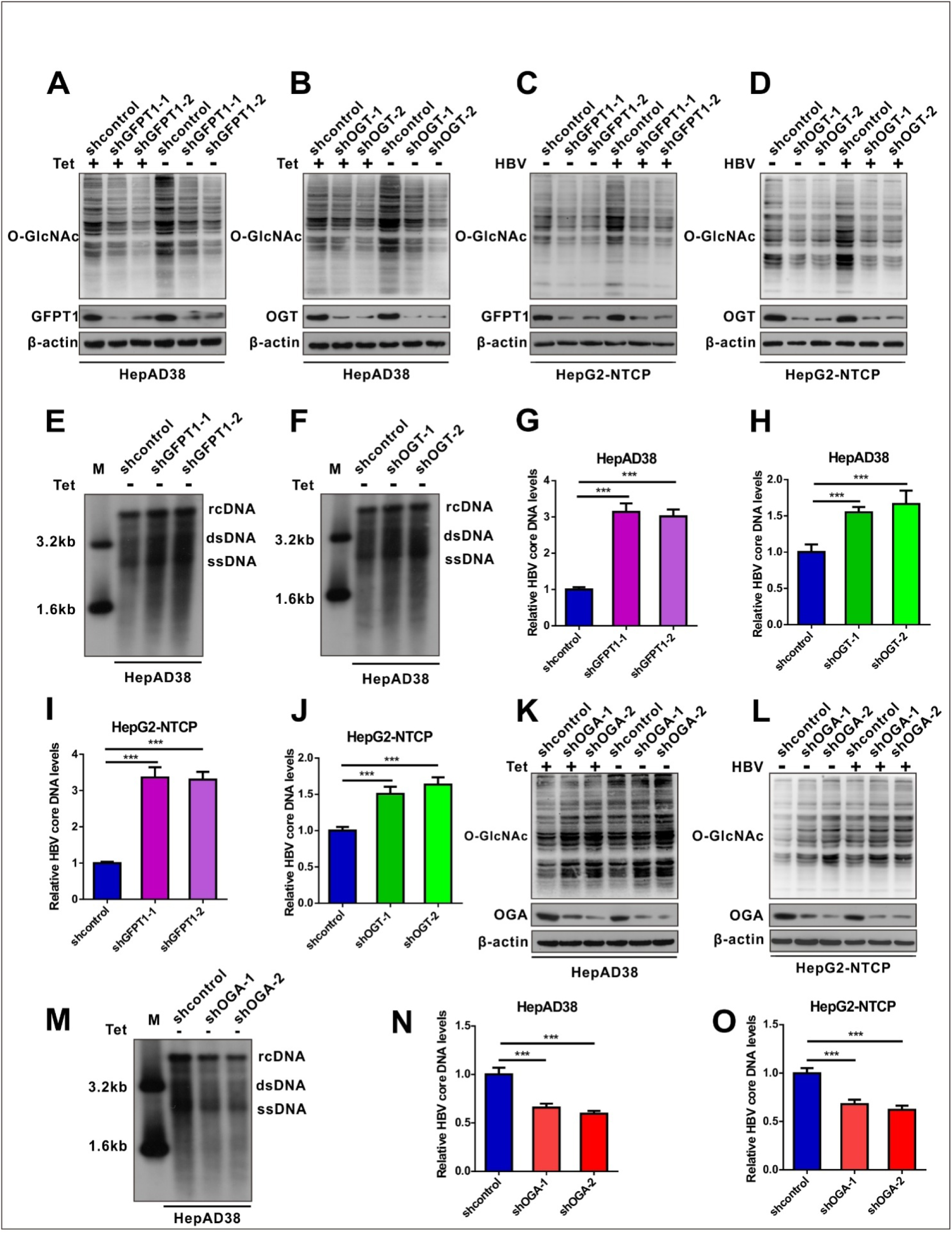
shRNA-mediated inhibition of protein O-GlcNAcylation enhances HBV replication. (**A**-**D**) Immunoblot of total O-GlcNAc from tetracycline-inducible HepAD38 cells (A-B) and HBV-infected HepG2-NTCP cells (C-D) following shRNA-mediated knockdown of GFPT1 and OGT. (**E**-**H**) Southern blot analysis of HBV DNA (E-F) and qPCR quantification of HBV core DNA levels (G-H) in stable HBV-expressing HepAD38 cells treated as above, n=9. (**I**-**J**) Quantification of HBV core DNA levels in HBV-infected HepG2-NTCP cells treated as indicated using qPCR, n=9. (**K**-**L**) Immunoblot of total O-GlcNAc from OGA-knockdown HepAD38 (Tet-off) cells (K) and OGA-knockdown HBV-infected HepG2-NTCP cells (L). (**M**) Southern blot analysis of HBV DNA in stable HBV-expressing HepAD38 cells treated as in K. (**N**-**O**) Quantification of HBV core DNA levels in stable HBV-expressing HepAD38 cells (N) and HBV-infected HepG2-NTCP cells (O) treated as in (M) using qPCR, n=9. Data are expressed as the mean ± SD. *P* values were derived from one-way ANOVA in G-H, I-J, and N-O; (\*\*\**P*< 0.001).

### OGT mediates O-GlcNAcylation of SAMHD1 upon HBV infection

To further investigate the mechanism by which OGT-mediated protein O-GlcNAcylation promotes host antiviral innate immunity during HBV infection, we screened putative O-GlcNAc-modified proteins in HepAD38 (Tet-off) cells using the immunoprecipitation assay coupled with mass spectrometry (IP-MS). Cell lysates were immunoprecipitated with O-GlcNAc antibodies and analyzed by LC-MS/MS. A total of 1,034 candidate O-GlcNAc-modified proteins were identified (Supplementary Table 1). Gene ontology analysis showed that several proteins were involved in innate immune and inflammatory responses (Supplementary Fig. 5A). We next focused on SAMHD1, which plays an important role in promoting host antiviral innate immunity (Ballana & Esté, 2015). Interactions between OGT and SAMHD1 were demonstrated by co-immunoprecipitation (co-IP) experiments in HepG2 cells (Fig. 4A-B). Confocal analysis indicated that OGT and SAMHD1 are co-localized in the nucleus (Fig. 4C). We subsequently constructed three SAMHD1 deletion mutants (Fig. 4D) and showed that the SAM domain of SAMHD1 is required for its interaction with OGT (Fig. 4E). Immunoprecipitated Flag-tagged SAMHD1 exhibited a strong O-GlcNAc modification signal in HEK293 cells upon treatment with the OGA inhibitor PUGNAc (Fig. 4F). Meanwhile, HBV replication enhanced SAMHD1 O-GlcNAcylation in HepAD38 (Tet-off) cells (Fig. 4G) and HBV-infected HepG2-NTCP cells (Supplementary Fig. 5B).These results were further confirmed by affinity chromatography using the succinylated wheat germ agglutinin (sWGA), a modified lectin that specifically binds O-GlcNAc-containing proteins (Fig. 4H-I). Collectively, these data indicate that SAMHD1 interacts with and can be O-GlcNAcylated by OGT upon HBV infection.

**Fig. 4.**
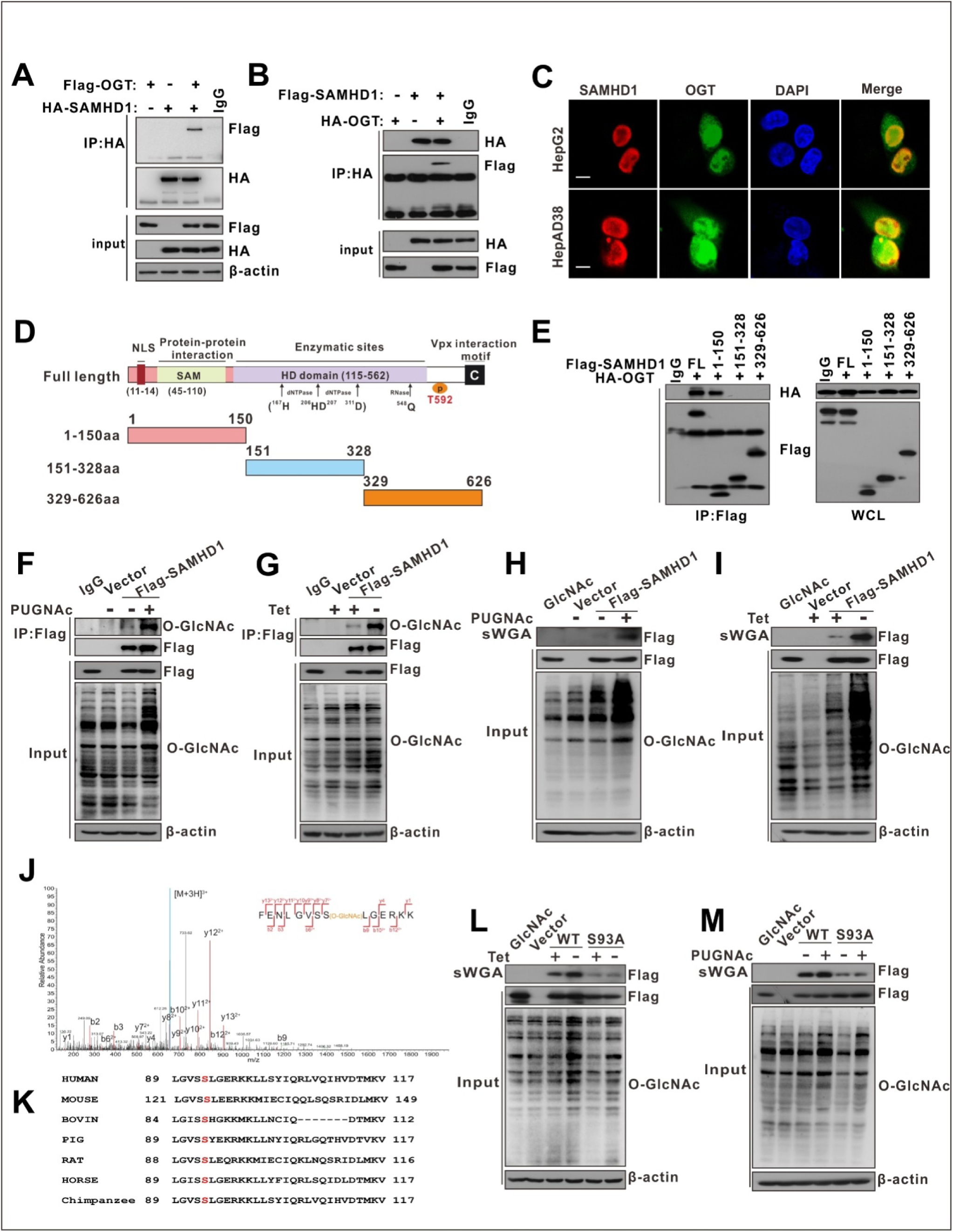
OGT mediates O-GlcNAcylation of SAMHD1 on Ser93. (**A**) Immunoprecipitation (IP) of SAMHD1 with anti-HA antibody in HEK293T cells co-transfected with Flag-OGT and HA-SAMHD1 expression constructs. The immunoprecipitated and input proteins were probed with the indicated antibodies. (**B**) Immunoprecipitation of OGT with anti-HA antibody in HEK293T cells co-transfected with HA-OGT and Flag-SAMHD1 expression constructs. (**C**) Representative confocal images of HepG2 (top) and HepAD38 cells (bottom) co-transfected with FLAG-SAMHD1 and HA-OGT. DAPI (blue) was used to counterstain nuclei. Scale bar, 10 μm. (**D**-**E**) Theinteraction between OGT and the full-length or the truncated SAMHD1 (1-150aa, 151-328aa, 329-626aa), as indicated in the diagram (D), were determined by Co-IP in HEK293T cells(E). (**F**) HEK293T cells were transfected with the Flag-SAMHD1 construct and the control vector for 48 h and treated with 100 μMPUGNAc for 12 h. Following cell lysis, SAMHD1 was immunoprecipitated using anti-FLAG M2 Agarose Beads. The immunoprecipitated and input proteins were probed with an anti-O-GlcNAc or anti-Flag antibody. (**G**) Immunoprecipitation of SAMHD1 with anti-Flag M2 agarose in tetracycline-inducible HepAD38 cells transfected with Flag-SAMHD1 and the control vector. (**H**-**J**) HEK293T cells (H) were treated as in (F) and tetracycline-inducible HepAD38 cells (I) were treated as in (G). After cell lysis, O-GlcNAc-modified proteins were purified using succinylated wheat germ agglutinin (sWGA)-conjugated agarose beads and probed with an anti-Flag or anti-O-GlcNAc antibody. GlcNAc served as a negative control. (**J**) LC-MS/MS analysis of FLAG-tagged SAMHD1 identified Ser93 as the SAMHD1 O-GlcNAcylation site. Tandem MS spectrum of the +2 ion at m/z 508.97 corresponding to O-GlcNAcylated SAMHD1 peptide FENLGVSSLGERKK is shown. (**K**) Multiple sequence alignment of SAMHD1 in different species. (**L**-**M**) SAMHD1-KO HepAD38 cells were transfected with empty vector, Flag-tagged SAMHD1 WT, or S93A mutant (l). HEK293T cells were transfected with the above plasmids described in (L) and treated with 100 μMPUGNAc for 12 h (M). Cell lysates were purified using sWGA-conjugated agarose beads and probed with an anti-Flag or anti-O-GlcNAc antibody.

### OGT-mediated O-GlcNAcylation on Ser93 enhances SAMHD1 stability

Next, we sought to map the O-GlcNAcylation site(s) on SAMHD1. Flag-tagged SAMHD1 was purified from HepG2-HBV1.3 cells and analyzed by MS. As shown in Fig. 4J, SAMHD1 was O-GlcNAcylated on Ser93 (S93). Interestingly, SAMHD1 S93 is well conserved among mammalian species (Fig. 4K). We then generated site-specific point mutants of SAMHD1. Mutation of S93 with Ala (S93A) largely reduced O-GlcNAc signal (Fig. 4L-M, and Supplementary Fig. 5C). To further examine the effect of O-GlcNAcylation on SAMHD1 stability, Flag-tagged wild-type or S93A mutant SAMHD1 was overexpressed alone or with shOGT in HepAD38 cells. The stability of exogenous SAMHD1 was decreased upon the expression of shOGT or S93A mutant (Fig. 5A-D). Moreover, SAMHD1 stability and ubiquitination was increased upon HBV infection (Fig. 5A-E). Furthermore, the administration of PUGNAC dramatically suppressed total and K48-linked ubiquitination of wild-type SAMHD1 (Fig. 5F); however, the effect on S93A ubiquitination was minimal (Fig. 5G). The S93A mutant was more ubiquitinated than wild-type SAMHD1 (Fig. 5G). These data indicate that O-GlcNAcylation of SAMDH1 at Ser93 stabilizes SAMHD1 by preventing its ubiquitination.

**Fig. 5.**
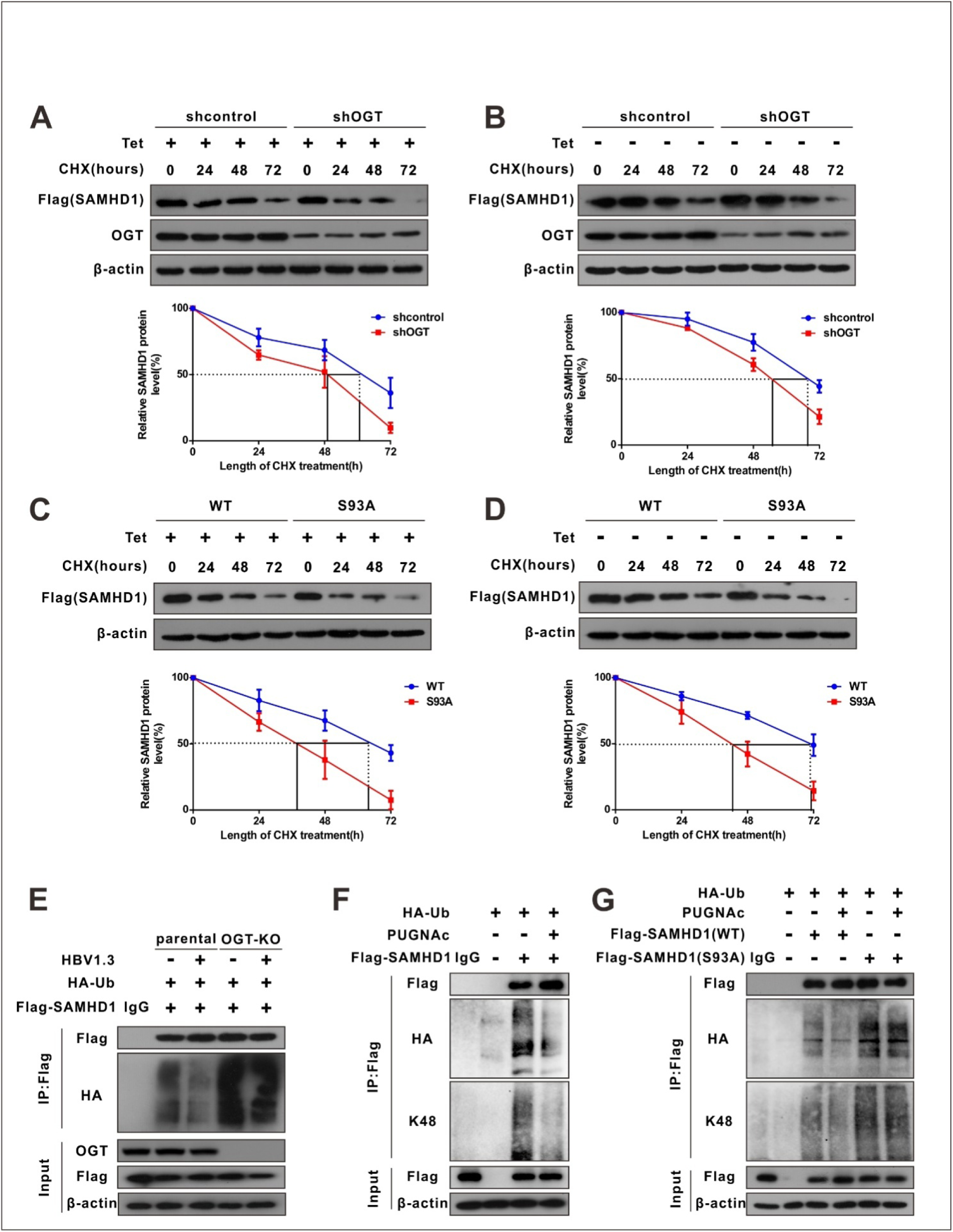
OGT-mediated O-GlcNAcylation on Ser93 enhances SAMHD1 stability. (**A**-**B**) Representative images of Flag-tagged SAMHD1 protein in non-infected or HBV infected SAMHD1 KO HepAD38 cells. Cells were transfected with Flag-tagged SAMHD1 and treated with 100 μM CHX for the indicated time.SAMHD1 band intensity was quantified using ImageJ,n=3. CHX, Cycloheximide. KO, knockout. (**C**-**D**) Immunoblots of SAMHD1. SAMHD1-KO HepAD38 cells treated with (Off) or without (On) tetracycline were transfected with Flag-tagged SAMHD1 WT or S93A mutant and treated with 100 μM CHX,n=3. (**E**) SAMHD1 ubiquitination in OGT-knockout HBV-infected HepG2 cells in the presence of HA-tagged ubiquitin. After cell lysis, SAMHD1 was immunoprecipitated using anti-FLAG M2 antibody. Immunoprecipitated and input proteins were probed with the indicated antibodies. (**F**-**G**) HEK293T cells were co-transfected with HA-Ub and Flag-SAMHD1 (F), Flag-tagged SAMHD1 WT or S93A mutant (G) and treated with 100 μMPUGNAc for 12 h. After cell lysis, SAMHD1 was immunoprecipitated using anti-FLAG M2 antibody. Immunoprecipitated and input proteins were probed with the indicated antibodies.

### O-GlcNAcylation of SAMHD1 on Ser93 enhances its antiviral activity

It is known that the tetramer conformation of SAMHD1 is required for its dNTP triphosphohydrolase (dNTPase) activity (Yan *et al*, 2013). Herein, we sought to determine whether the S93A mutant affects SAMHD1 tetramerization and dNTPase activity. Recombinant WT and S93A SAMHD1 were expressed and purified (Supplementary Fig. 6A-B). We found that S93A mutation destabilized SAMDH1 tetramers in HepAD38 cells (Fig. 6A) and reduced its dNTPase activity *in vitro* (Supplementary Fig. 6C-D). To test the effect of S93 O-GlcNAcylation on SAMHD1 antiviral activity, we deleted endogenous SAMHD1 in our HBV cell models and THP-1 cells using CRISPR-Cas9-mediated gene editing, and transfected wild-type or SAMHD1 variants into SAMHD1-knockout HepAD38 (Tet-off) (Fig. 6B), AdHBV-1.3-infected HepG2 (Fig. 6C), and HepG2-NTCP cells. A phospho-mimetic mutation (T592E) was used as a control that also decreased SAMHD1 dNTPase activity and abrogated its antiviral activity (Sommer *et al*, 2016). Both southern blotting (Fig. 6B-C) and qPCR (Fig. 6D-F) results indicated that S93A mutation impairs the ability of SAMHD1 to inhibit HBV replication *in vitro*. A previous study showed that SAMHD1 dNTPase activity is essential for HIV-1 restriction (Hansen *et al*, 2014). Therefore, we investigated the effect of SAMHD1 O-GlcNAcylation on HIV-1 infection. THP-1 cells were infected with a vesicular stomatitis virus G (VSV-G) protein pseudotyped HIV-1 molecular clone carrying the luciferase gene reporter, and virus replication was assessed by quantifying luciferase activity. Our results showed that protein O-GlcNAcylation was increased upon HIV-1 infection in THP-1 cells (Fig. 6G). Subsequently, wild-type or SAMHD1 variants were transfected into SAMHD1-KO THP-1 cells. S93A mutation also impaired the ability of SAMHD1 to restrict HIV-1 replication in this single-round HIV-1 infection model (Fig. 6H). Treatment of cells with the GFPT inhibitor 6-diazo-5-oxo-L-norleucine (DON) and the OGT inhibitor ST045849 significantly increased luciferase activity, whereas treatment with the OGA inhibitor PUGNAc reduced luciferase activity (Fig. 6I). Taken together, these results indicate that O-GlcNAcylation of SAMHD1 S93 promotes its antiviral activity *in vitro*.

**Fig. 6.**
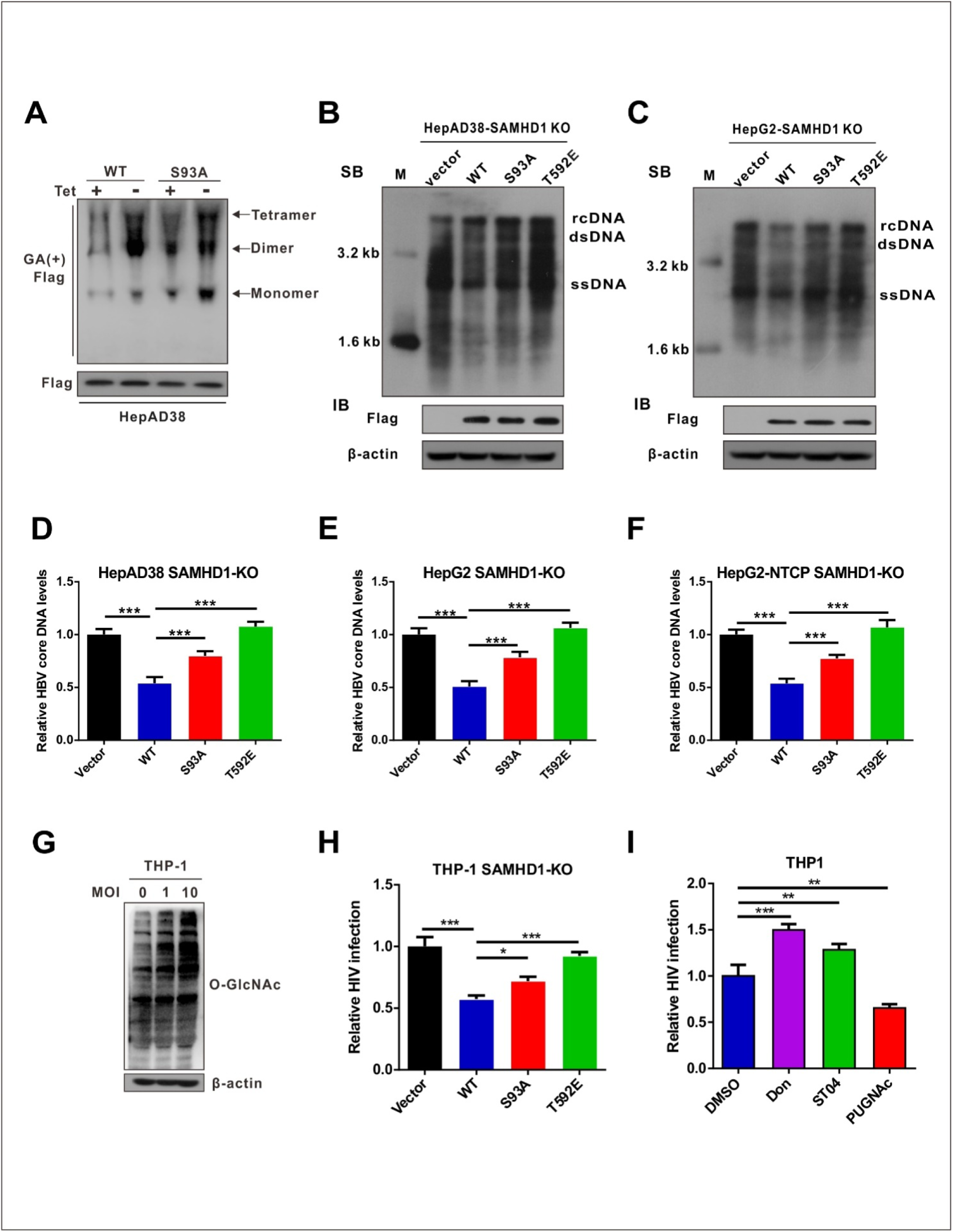
O-GlcNAcylation of SAMHD1 on Ser93 is important for its antiviral activity. (**A**) Changes in the oligomeric state of SAMHD1 upon HBV infection. SAMHD1-KO HepAD38 cells with tetracycline inducible (Tet-off) HBV expression were transfected with the Flag-tagged SAMHD1 WT or S93A mutant construct. Cells were treated with glutaraldehyde (GA) and whole-cell lysates were probed with an anti-Flag antibody. (**B**-**C**) HepAD38 cells with stable HBV-expressing (B) and HBV-infected SAMHD1-KO HepG2 cells (C) were transfected with Flag-tagged SAMHD1 WT, S93A mutant, or T592E mutant. HBV DNA levels were determined by southern blot analysis. (**D**-**F**) SAMHD1-KO HepAD38 cells with stable HBV-expressing (D), HBV-infected SAMHD1-KO HepG2 (E) and SAMHD1-KO HepG2-NTCP cells (F) were transfected with the above plasmids described in (B). HBV core DNA levels were determined by qPCR. n=9. (**G**) SAMHD1 KO-THP-1 cells were differentiated overnight with PMA (100 μM) before infecting with HIV-LUC-G (MOI=0, 1, or 10) for 48 h. Thereafter, the cells were lysed and total O-GlcNAc levels were determined by western blotting. β-actin was used as a loading control. (**H**) SAMHD1 KO-THP-1 cells were differentiated overnight and infected with HIV-LUC-G (MOI=1) for 24 h. Thereafter, they were transfected with Flag-tagged SAMHD1 WT, S93A mutant, or T592E mutant for 48 h. Luciferase activity was measured and normalized for protein concentration. n=3. (**I**) SAMHD1 KO-THP-1 cells were differentiated overnight and infected with HIV-LUC-G (MOI=1) for 24 h. Cells were then treated with Don (30 μM, 24 h), ST04 (100 μM, 24 h), or PUGNAc (100 μM, 48 h), and luciferase activity was measured. n=3. Data are expressed as the mean ± SD. *P* values were derived from one-way ANOVA in D-F, H-I. (* *P*<0.05, ** *P*<0.01, \*\*\**P*<0.001).

### HBV infection promotes UDP-GlcNAc biosynthesis and O-GlcNAcylation *in vivo*

We used an HBV-transgenic (HBV-Tg) mouse model to verify our results *in vivo* (Fig.7A).The level of O-GlcNAcylation was significantly higher in the liver tissues of HBV-Tg mice than in those of normal C57BL/6 mice (Fig. 7B). Consistent with our *in vitro* data, the administration of DON significantly reduced UDP-GlcNAc levels (Fig. 7C) and stimulated HBV replication (Fig. 7D-F) in the mouse model of HBV infection, whereas the administration of Thiamet G decreased serum HBV DNA (Fig. 7E), liver HBcAg (Fig. 7F) and HBV DNA (Fig. 7G) levels in mice. Protein O-GlcNAcylation levels in the liver tissues of HBV-Tg mice were increased upon Thiamet G administration, but decreased upon DON administration (Fig. 7H). These results indicate that Thiamet G can promote host antiviral immunity by increasing protein O-GlcNAcylation. Finally, we examined UDP-GlcNAc biosynthesis and O-GlcNAcylation levels in patients with chronic hepatitis B (CHB). The levels of serum UDP-GlcNAc (Fig. 7I), GLUT1 protein (Fig. 7J), and total O-GlcNAcylation (Fig. 7J and 7K) were markedly higher in the liver tissues of patients with CHB than in those of normal controls. In addition, SAMHD1 O-GlcNAcylation was significantly increased in the liver tissues of the patients with CHB (Fig. 7K). Overall, our study suggests that HBV infection upregulates GLUT1 expression and increases UDP-GlcNAc biosynthesis and O-GlcNAcylation *in vivo*. As an essential O-GlcNAcylated protein, SAMHD1 can exert its antiviral activity and elicit a robust host innate immune response against HBV infection.

**Fig. 7.**
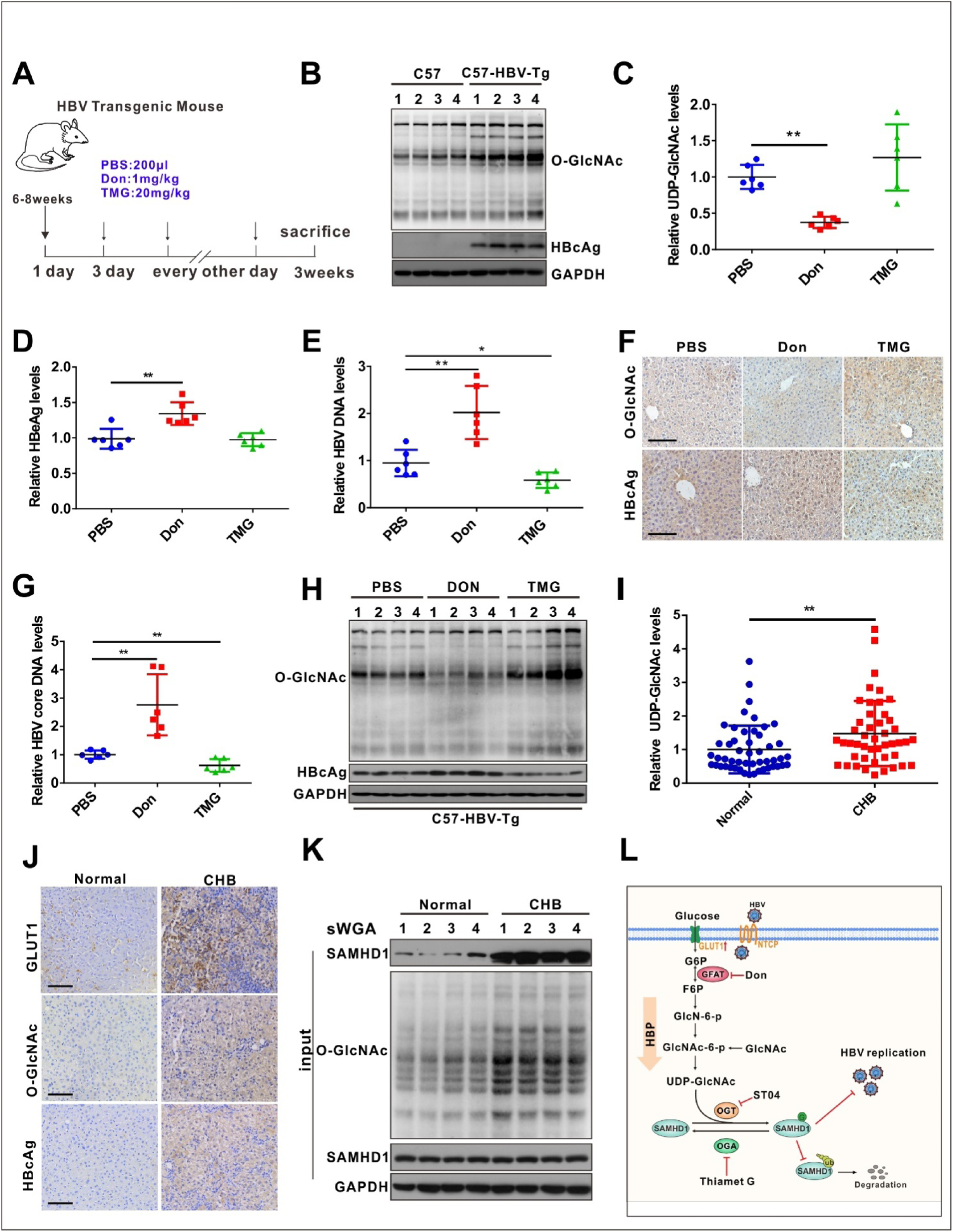
HBV infection promotes UDP-GlcNAc biosynthesis and protein O-GlcNAcylation *in vivo*. (**A**) Six- to eight-week-old HBV transgenic mice were intraperitonally injected with Don (1 mg/kg body weight) and TMG (20 mg/kg body weight) or PBS (control) every other day for 10 times. The mice were sacrificed on day 20 post-treatment. (**B**) Immunoblotting of total O-GlcNAc in HBV transgenic mice. (**C**) Fold change in the expression of UDP-GlcNAcin mouse liver tissues was determined by UHPLC-QTOF-MS. n=6 per group. (**D**-**E**) Serum HBeAg and HBV DNA levels in mice. n=6 per group. (**F**) O-GlcNAc and HBcAg detection in mouse liver tissues, Scale bar, 50 μm. (**G**) Quantification of HBV core DNA levels in mouse liver tissues using qPCR. n=6. (**H**) Immunoblot of total O-GlcNAc in HBV transgenic mice treated as in (A). (**I**) Fold change in the expression of UDP-GlcNAcin the liver tissues of patients with CHB was determined by UHPLC-QTOF-MS. (Normal=50, CHB=46). (**J**) GLUT1, O-GlcNAc, and HBcAg detection in liver tissue specimens from patients with CHB. Scale bar, 50 μm. (**K**) Liver tissue lysates from patients with CHB were purified using sWGA-conjugated agarose beads and probed with an anti-SAMHD1 or anti-O-GlcNAc antibody. (**L**) Proposed working model of this study. Data are expressed as the mean ± SD. *P* values were derived from one-way ANOVA in C-E, G, and from unpaired, two-tailed Student’s *t*-test in I. (* *P*<0.05, ** *P*< 0.01).

## Discussion

Although previous studies have demonstrated that HBV infection can alter glucose metabolism in host cells, the role and underlying mechanisms of metabolic regulation of antiviral immune responses remain elusive. In this study, we demonstrate that HBV increases GLUT1 expression on hepatocyte surface, thereby facilitating glucose uptake. This enhanced nutrient state consequently provides substrates to HBP to produce UDP-GlcNAc, leading to an increase in protein O-GlcNAcylation. Importantly, we found that pharmacological or transcriptional inhibition of HBP and O-GlcNAcylation can promote HBV replication. Furthermore, we showed that OGT-mediated O-GlcNAcylation of SAMHD1 on Ser93 is critical for its antiviral activity. Our results therefore indicate that O-GlcNAcylation can positively regulate host antiviral immune response against HBV infection.

Similar to the metabolic reprogramming in proliferating cancer cells, virus reprogram host cell metabolism. It has been reported that several viruses increase glucose consumption and reprogram glucose metabolism in the host cell (Purdy & Luftig, 2019; Thaker *et al*, 2019). GLUT1 expression was increased in host cells infected with HIV-1 (Loisel-Meyer *et al*, 2012; Palmer *et al*, 2014), Kaposi’s sarcoma-associated herpes virus (Gonnella *et al*, 2013), dengue virus (Fontaine *et al*, 2015), and Epstein-Barr virus (Zhang *et al*, 2017). Our findings are consistent with previous transcriptome-wide analyses, which have also shown HBV-mediated upregulation of GLUT1 (Lamontagne *et al*, 2016). It has been suggested that HBV pre-S2 mutant increases GLUT1 expression via mammalian target of rapamycin signaling cascade, leading to enhanced glucose uptake (Teng *et al*, 2015, 2). However, the precise molecular mechanism by which HBV upregulates GLUT1 remains poorly understood.

The enhanced glucose uptake by glucose transporter not only accelerates glycolysis, but may also increase flux into branch pathways, such as the pentose phosphate pathway and HBP, which occur in cancer cells (Ma & Vosseller, 2014). Previous studies have reported that HBP plays an important role in host innate immunity. Consistent with the results of a previous study with HepG2.2.15 cells (Li *et al*, 2015), our results showed that HBV infection can promote HBP activity and increase UDP-GlcNAc levels in different cell models. Li *et al.* reported that enhanced HBP activity is essential for HBV replication because pharmacological or transcription suppression of *GFPT1* inhibits HBV replication in HepG2.2.15 cells. However, they did not use an *in vivo* HBV model to study the underlying mechanism. In contrast, we showed that blockade of HBP promotes HBV replication, whereas stimulation of HBP significantly suppresses HBV replication both *in vitro* and *in vivo*. In addition, we observed similar results upon HIV-1 infection using a single-round infection model. Although we could not exclude the possibility that differences between HBV cell models cause this discrepancy, our results show that increased HBP flux and hyper-O-GlcNAcylation can upregulate host antiviral innate response. Several other studies have reported that HBP and/or protein O-GlcNAcylation promotes host antiviral immunity against RNA viruses, including VSV (Li *et al*, 2018), influenza virus (Song *et al*, 2019), and hepatitis C virus (Herzog *et al*, 2019). Thus, the present study confirms and expands our current understanding of the antiviral activity of HBP and protein O-GlcNAcylation upon DNA virus infection, which is similar to its antiviral activity upon infection by certain RNA viruses.

By characterizing the role of protein O-GlcNAcylation during HBV replication, we uncovered SAMHD1 as an important target of OGT and established a link between O-GlcNAcylation and antiviral immune response against HBV infection. SAMHD1, an effector of innate immunity, can restrict most retroviruses (such as HIV-1) and several DNA viruses (including HBV) by depleting the intracellular pool of dNTPs (Ballana & Esté, 2015). Several post-translational modifications, including phosphorylation (White *et al*, 2013, 1) and ubiquitination (Li *et al*, 2019b) have been reported to be critical for SAMHD1 function. Herein, we identified Ser93 as a key O-GlcNAcylation site on SAMHD1 using LC-MS/MS. Importantly, loss of O-GlcNAcylation by S93A mutation increased K48-linked ubiquitination, thus decreased the stability and dNTPase activity of SAMHD1, suggesting that O-GlcNAcylation promotes the antiviral activity of SAMHD1.

Because these results demonstrated the importance of protein O-GlcNAcylation in host antiviral innate immunity against HBV, we proposed that an increase in SAMHD1 O-GlcNAcylation by inhibiting OGA activity could be used as a potential antiviral strategy. This is in line with recent results indicating that increased MAVS O-GlcNAcylation is essential to activate host innate immunity against RNA viruses (Li *et al*, 2018; Song *et al*, 2019). However, hyper-O-GlcNAcylation has been reported to stabilize several oncogenic factors in several cancers associated with oncogenic virus infection (Makwana *et al*, 2019). Human papillomavirus 16 E6 protein can upregulate OGT and stabilize c-MYC via O-GlcNAcylation, thus promoting HPV-induced carcinogenesis (Zeng *et al*, 2016). Herzog *et al*. demonstrated that protein O-GlcNAcylation is involved in HCV-induced disease progression and carcinogenesis (Herzog *et al*, 2019). Thus, the role of protein O-GlcNAcylation in HBV pathogenesis and the antiviral response through enhanced protein O-GlcNAcylation remain to be further studied.

In conclusion, we uncovered a link between metabolic reprogramming and antiviral innate immunity against HBV infection. We demonstrated that HBV infection upregulates GLUT1 expression and promotes HBP flux *in vitro* and *in vivo*. In addition, increased UDP-GlcNAc biosynthesis and hyper-O-GlcNAcylation can enhance host antiviral innate response. Mechanistically, OGT-mediated O-GlcNAcylation of SAMHD1 on Ser93 stabilizes SAMHD1 and enhances its antiviral activity (Fig. 7l). This study broadens our understanding of SAMHD1 post-translational modification and provides new insights into the importance of HBP and protein O-GlcNAcylation in antiviral innate immunity.

## Materials and Methods

### Animal models

HBV-transgenic (HBV-Tg) mice (n = 6 for each group) were kindly provided by Prof. Ning-shao Xia, School of Public Health, Xiamen University(Huang *et al*, 2006). C57BL/6J mice (6- to-8-week-old, six per group) were provided by the Laboratory Animal Center of Chongqing Medical University (SCXK (YU) 2018-0003). Mice were intraperitoneally injected with Don (1 mg/kg body weight), Thiamet G (20 mg/kg body weight), or PBS (control) every other day for 10 times. On day 20 post-administration, mouse serum and liver tissue specimens were collected for real-time PCR, southern blotting, and immunohistochemical staining. Mice were treated in accordance with the guidelines established by the Institutional Animal Care and Use Committee at the Laboratory Animal Center of Chongqing Medical University. The animal care and use protocols adhered to the National Regulations for the Administration of Laboratory Animals to ensure minimal suffering.

### Samples from patients with chronic hepatitis B virus infection

The study protocol was approved by the Medical Ethics Committee of Chongqing Medical University. Informed consent was obtained from patients who met the inclusion criteria for chronic HBV infection.

### Metabolites analysis

To extract metabolites from quenched serum/plasma samples or cell culture supernatants, 400 μL chilled methanol: acetonitrile (2:2, v/v) was added to 100 μL of each sample. The mixture was vortexed three times for 1 min each with 5-min incubation at 4°C after each vortexing step. After the final vortexing step of 30 s, the mixture was incubated on ice for 10 min. Thereafter, 100 μL chilled HPLC-certified water was added to the samples, mixed for 1 min, and centrifuged at 13,000g for 10 min at 4°C. Finally, the liquid phase (supernatant) of each sample was transferred into a new tube for UHPLC-QTOF-MS analysis in Shanghai Applied Protein Technology Co., Ltd. UDP-GlcNAc and glucose were quantified using targeted liquid chromatography-tandem mass spectrometry (LC-MS/MS). The data acquisition, principal component analysis, heatmap and pathway impact analysis were performed by Shanghai Applied Protein Technology Co., Ltd.

### Immunoprecipitation assay coupled with mass spectrometry (IP-MS)

HepAD38 (Tet-off) cell lysates were incubated overnight with an anti-O-GlcNAc antibody at 4°C, followed by a 4-h incubation with protein A/G agarose beads. Immunoprecipitated complexes were eluted and stained with Coomassie blue. Stained protein bands were sent to Shanghai Applied Protein Technology Co., Ltd for identification of potential O-GlcNAc-modified proteins. Protein bands were dissolved in 1 mL chilled methanol: acetonitrile: H_2_O (2:2:1, v/v/v) and sonicated at low temperature (30 min); this process was repeated twice. The supernatant was dried in a vacuum centrifuge. For LC-MS analysis, samples were re-dissolved in 100 μL acetonitrile: water (1:1, v/v). Sample analyses were performed using a UHPLC system (1290 Infinity LC, Agilent Technologies) coupled to a quadrupole time-of-flight analyzer (AB Sciex Triple TOF6600) at Shanghai Applied Protein Technology Co., Ltd.

### SAMHD1 O-GlcNAcylation site mapping

Mass spectrometry was performed to identify SAMHD1 O-GlcNAcylation sites, as described previously (Peng *et al*, 2017). Briefly, immunoprecipitated SAMHD1 from HEK293T cells was subjected to SDS-PAGE. The band corresponding to SAMHD1 was excised, digested overnight with trypsin, and subjected to liquid chromatography-tandem mass spectrometry (LC-MS/MS) analysis. An online LC-MS/MS setup consisting of an Easy-nLC system and an Orbitrap Fusion Lumos Tribrid mass spectrometer (Thermo Scientific, Germany) equipped with a nanoelectrospray ion source was used for all LC-MS/MS experiments. Raw MS files were searched against the UniProt database using MaxQuant software (version 1.5.2.8). The fixed modification was set to C (carbamidomethyl) and the variable modifications were set to M (oxidation), protein N-term (acetyl), and S/T (O-GlcNAc). The peptide tolerance for the first search was set at 20 ppm and that for the main search was set at 6 ppm. The MS/MS tolerance was 0.02 Da. The false discovery level in PSM and protein was 1%. The match between runs was used and the minimum score for modified peptides was set at 40.

### Statistical Analysis

All data are expressed as the mean ± standard deviation (SD). All statistical analyses were performed using GraphPad Prism 5.0 software (GraphPad Software Inc.). Statistical significance was determined using one-way ANOVA for multiple comparisons. Student’s *t*-test was used to compare two groups. *P*<0.05 was considered statistically significant.

For detailed descriptions of other methods, please refer to **Supplementary Methods**.

## Acknowledgements

We are grateful to Dr. T.-C He (University of Chicago, USA) for providing the pAdEasy plasmid. We thank Prof. Bing Sun (Shanghai Institute of Biochemistry and Cell Biology, China) for providing the pLL3.7 vector. We also thank Prof. Cheguo Cai (Wuhan University, China) for providing the pNL4-3.Luc.R-E-plasmid. This work was supported byNational Natural Science Foundation of China (grant nos. 81872270, 81572683, 81661148057), the Natural Science Foundation Project of Chongqing (cstc2018jcyjAX0254, cstc2019jcyj-msxmX0587), the Major National S&T program (2017ZX10202203-004), the National Key Research and Development Program of China (2018YFE0107500), the Leading Talent Program of CQ CSTC (CSTCCXLJRC201719), and the Scientific Research Innovation Project for Postgraduate in Chongqing (grant nosCYB19168, CYS19193)

## Authors Contributions

NT, AH, and KW conceived the study and designed the experiments. JH, QG, YY and XJ performed most experiments and analyzed the data. WZ and LC performed SAMHD1 O-GlcNAcylation site mapping. YC and ZZ collected clinical samples. LL generated SAMHD1 mutants. QL assisted with HepG2-NTCP cell culture. YH, HZ and XLprovided guidance and advice. JH, QG, KW, and NT wrote the manuscript with all authors providing feedback.

